# Positive and Negative Retinotopic Codes in the Human Hippocampus

**DOI:** 10.1101/2024.09.27.615397

**Authors:** Peter A. Angeli, Adam Steel, Caroline E. Robertson

## Abstract

The hippocampus sits at the apex of the visual hierarchy, yet little is known about the visual properties of this core memory structure. Recent work suggests that a latent, bivalent retinotopic code persists in large-scale memory networks at the cortical apex, scaffolding interactions with sensory networks. Here, we tested whether a bivalent retinotopic code also persists within the hippocampus. To do this, we leveraged high-resolution 7T functional MRI along with voxel-scale visual population receptive field (pRF) modeling in 7 densely-sampled individuals (5 female). Our findings reveal a robust, voxel-scale retinotopic code broadly distributed across subfields and along the long axis of the human hippocampus, comprised of roughly equal proportions of pRFs with positive and negative amplitude responses to visual stimulation. Hippocampal pRFs displayed canonical visual properties, including stable valence and visual field preferences across runs and a contralateral bias. Retinotopic structure also persisted at rest: hippocampal voxels with similar pRF locations were more strongly correlated than voxels representing different visual field locations. Finally, across the ventral visual stream, the prevalence of negative-amplitude pRFs increased with mnemonic involvement, culminating in the balanced, bivalent organization within the hippocampus. These findings support the view that sensory and mnemonic systems are coupled through a shared retinotopic code at the apex of the visual hierarchy.

**Significance Statement:** The hippocampus is closely coupled to the visual system, and recent work has challenged the classical view that visual coding schemes, like retinotopy, do not persist into the hippocampus. Here, we use high-resolution precision fMRI to robustly characterize a bivalent retinotopic code in the human hippocampus, consisting of both typical positive and atypical negative responses. We further show that this code predicts functional connectivity within the hippocampus even during non-visual tasks, suggesting that this bivalent retinotopic code may reflect an intrinsic organizational principle of the hippocampus relevant for perceptual-mnemonic segregation and integration.

## Introduction

The hippocampus serves as a critical hub for sensory-mnemonic interactions, integrating sensory information with memory processing to support the encoding and retrieval of episodic memories (Scoville and Milner, 1957; Cohen and Squire, 1980; Cohen and Eichenbaum, 1993). As such, hippocampal representations essentially rely on input from sensory systems, including the visual system (Vinogradova, 2001; Jeffery, 2007; Pereira et al., 2007). Yet relatively little is known about the visual coding properties of this core memory structure.

Classic models of cortical organization place the hippocampus at the apex of the visual hierarchy (Felleman and Van Essen, 1991) where it is thought to construct abstract, internal models of the external world. Canonically, hippocampal coding schemes are thought to be primarily allocentric and context-dependent (Scoville and Milner, 1957; O’Keefe and Dostrovsky, 1971; Cohen and Squire, 1980; Cohen and Eichenbaum, 1993; Eichenbaum and Cohen, 2014; Whittington et al., 2020), fundamentally distinct from sensory-driven retinotopic codes like those found in early visual areas (Sereno and Allman, 1991; Tootell et al., 1998). However, recent findings challenge this view. Work in rodents and nonhuman primates has shown that a significant proportion of hippocampal neurons exhibit egocentric visual coding (Acharya et al., 2016; Gulli et al., 2020; Purandare et al., 2022), and human fMRI research has identified visual field preferences in hippocampal voxels (Knapen, 2021; Silson et al., 2021). Thus, contrary to the classic view, these cross-species findings suggest the hippocampus may demonstrate egocentric visual coding, but evidence from cerebral cortex indicates these descriptions may be incomplete.

Recent studies of retinotopic coding in memory-related cortical regions, like the Default Network (DN), have observed atypical retinotopic coding alongside classic mnemonic responses. Specifically, many of the DN’s retinotopic responses exhibit a negative valence — that is, decreasing activity in response to visual presentation (Szinte and Knapen, 2020; Klink et al., 2021; Knapen, 2021; Steel et al., 2024b) — in contrast to the typical positive valence found in visual areas. This bivalent (positive and negative) retinotopic code structures a voxel-scale push-pull dynamic between perceptual and mnemonic cortical regions and networks (Steel et al., 2024b, 2024a). One hypothesis is that a similar bivalent retinotopic code might exist in the hippocampus, given the structure’s well-characterized mnemonic properties and connectivity with the DN (Greicius et al., 2004; Vincent et al., 2006; Kahn et al., 2008).

In this study, we used high-resolution 7T functional MRI from visual mapping tasks to test whether a bivalent retinotopic code is present within the human hippocampus. In brief, our results show that the human hippocampus contains a robust retinotopic code, with a balanced proportion of positive and negative population receptive fields (pRFs), evenly distributed throughout the hippocampus. Importantly, this code is reliable: pRF amplitude and valence are consistent across runs of the retinotopic mapping task. Both positive and negative pRFs demonstrate a bias towards the contralateral visual field, and we found the proportion of negative pRFs increases moving up the ventral visual stream hierarchy, parallelling mnemonic involvement. Finally, during resting-state fixation, retinotopic preferences strongly influence the correlation of activity among hippocampal pRFs, suggesting that this bivalent retinotopic code may be broadly relevant for mnemonic information processing in the hippocampus.

## Methods

### Overview

We used data made publicly available through the Natural Scenes Dataset (NSD, Allen et al., 2022) that included 8 participants scanned extensively at 7T over the course of several days.

The high resolution of the data allowed for detailed explorations of hippocampal activity. Relevant to the present study, each participant completed *both* three runs of sweeping bar retinotopic mapping task and at least 14 runs of a resting-state fixation task (range: 14 – 34 runs). Retinotopy and fixation data were minimally preprocessed using manual ICA denoising, and all subsequent analyses were performed in participant-specific volume space. To allow for the identification of negative retinotopic responses, population receptive field (pRF) modeling was performed using AFNI’s mapping procedure (3dNLfim). We did not regress global signal, and data were left unsmoothed.

### Data Preparation

#### Participants and Data Acquisition

Data from the eight participants (ages 19 – 32) included in the NSD release were used for the current study. One participant (subj03) was excluded from all analyses due to an insufficient quantity of resting-state fixation data that passed our quality control criteria (see Quality Control and Preprocessing below). The final set of seven participants had a mean age of 26.1 yr (7 right-handed; 5 women) and had normal or corrected-to-normal visual acuity. All participants provided informed consent for the study.

MRI data was acquired at the Center for Magnetic Resonance Research at the University of Minnesota as described in Allen et al. (2022), and the study was approved by the University of Minnesota institutional review board. Briefly, anatomical scans were acquired using a 3T Siemens Prisma scanner with a 32-channel head coil, including several T_1_-weighted magnetization prepared rapid acquisition gradient echo (MPRAGE, voxel size = 0.8 mm^3^ isotropic, TR = 2,400 ms, TE = 2.22 ms, flip angle = 8°, in-plane acceleration factor = 2) and T_2_-weighted sampling perfection with application-optimized contrasts using different flip angle evolution sequence (SPACE, voxel size = 0.8 mm^3^ isotropic, TR = 3,200 ms, TE = 563 ms, in-plane acceleration factor = 2) acquisitions. The remaining data was acquired on a 7T Siemens Magnatom scanner and a single-channel-transmit, 32-channel-receive head coil. Functional data were gradient-echo echo-planar scans (voxel size = 1.8 mm^3^ isotropic, TR = 1,600 ms, TE = 22 ms, flip angle = 62°, matrix = 120 x 120, field-of-view = 216 mm x 216 mm), corrected with dual-echo fieldmaps (matched slice prescription to EPI, voxel size = 2.2 mm x 2.2 mm x 3.6 mm TR = 510 ms, TE = 8.16 ms, TE = 9.18 ms). For improved hippocampal segmentation, an additional T_2_-weighted turbo-spin echo scan was collected (voxel size = 0.357 mm x 0.357 mm x 1.5 mm, 56 oblique slices perpendicular to anterior-posterior axis of the hippocampus, TR = 16,000 ms, TE = 53 ms, in-plane acceleration factor = 2).

#### Task Paradigms

The retinotopic mapping paradigm (adapted from the Human Connectome Project Retinotopy Dataset; Benson et al., 2018, RETBAR1 and RETBAR2) consisted of a bar aperture slowly moving across the visual field, revealing objects at multiple scales (i.e. faces, houses, and objects) superimposed on a pink-noise background (Fig. 1A). Both the apertures and textures updated at a rate of 15 Hz, and covered a circular region with a diameter of 8.4°. For each 300 second run, participants were instructed to fixate on a small (0.2° x 0.2°) semi-transparent dot in the center of the screen, and indicate with a button press whenever the color of the dot changed (which occurred every 1 – 5 sec). Three runs of this task were collected, for a total of 15 minutes of data. Additional wedge/ring runs were also acquired as part of the NSD, but consistent with other studies characterizing −pRFs (Szinte and Knapen, 2020; Klink et al., 2021; Steel et al., 2024b) here we consider only the bar stimulus.

**Fig. 1.**
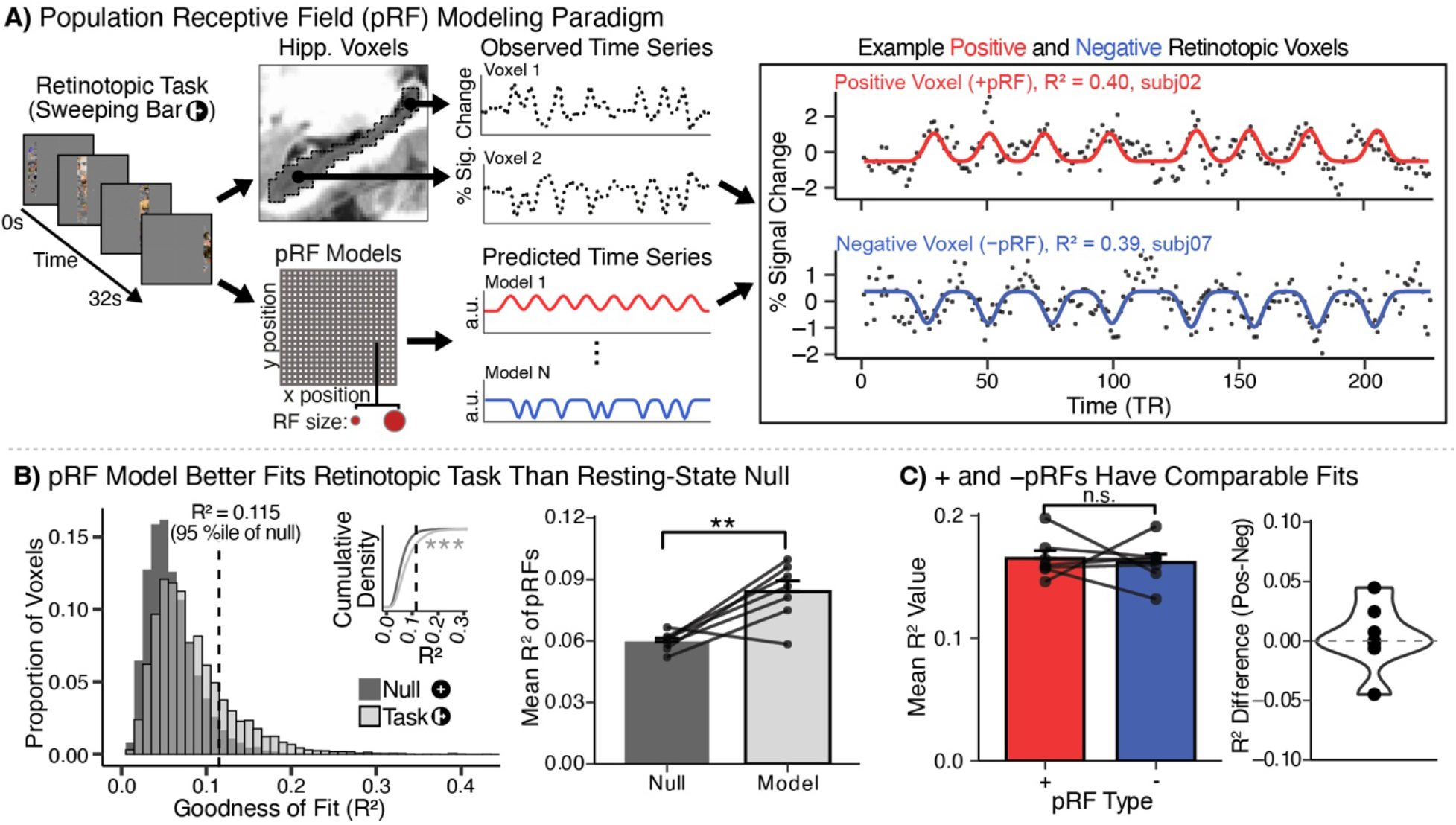
Reliable positive and negative population receptive fields (pRF) in the human hippocampus. **A. Schematic of fMRI population receptive field (pRF) modeling procedure.** Voxel time series from a sweeping bar visual mapping task (Benson et al., 2018; Allen et al., 2022) were fit with a 2D Gaussian pRF model varying in center location (x,y), size (degrees visual angle), and amplitude (positive or negative). Voxels with a positive amplitude parameter are termed +pRFs, while those with negative amplitude parameters are termed −pRFs. Schematic of theoretical time courses (points) with model prediction (line) highlight the impact of amplitude. Example time courses (right) illustrate stimulus-evoked activation for +pRFs and suppression for −pRFs relative to baseline. **B. Task-derived pRF model fits in the hippocampus exceeded those obtained from a null distribution generated using resting-state fixation data.** Across hippocampal voxels (pooled across n=7 participants), task-derived model fits were significantly right-shifted relative to null fits. **C. Positive and negative pRFs had similar goodness-of-fit (R^2^) values after thresholding.** (Left) Subject-level mean R² for +pRFs (red) and −pRFs (blue). There was no significant difference between valences. (Right) Subject-level difference in mean R² between +pRFs and −pRFs (no significant difference). ***p < 0.001 **p < 0.01 *p < 0.05

In all resting-state fixation tasks, participants were instructed to fixate on a small white cross (0.5° x 0.5°) in the center of a gray screen. In each session where fixation tasks were acquired, one run was collected at the beginning of the session, and the other at the end. The first fixation run included no additional instructions. For the second fixation run, participants were instructed to take a deep breath 12 sec after the beginning of the run, when the fixation cross turned red for 4 sec. Each run of fixation was 300 sec, and between 14 and 34 runs (70-170 minutes of data) were collected for each participant.

#### Quality Control and Preprocessing

All data was assessed for quality prior to inclusion in the dataset as described in the original data release (Allen et al., 2022). Motivated by the sensitivity of resting-state functional connectivity to data quality, particularly motion, we performed additional quality control measures on the fixation data. We assessed the quality of resting-state fixation data using AFNI’s quality control report creation tool (APQC, Taylor et al., 2024), and excluded any fixation run with a maximum displacement of greater than 1.8 mm, or a framewise displacement greater than 0.12 mm. Runs with a maximum displacement between 1 mm and 1.8 mm, and a framewise displacement of between 0.1 mm and 0.12 mm, were assessed on a case-by-case basis, and excluded if the movement was the result of large shifts (approach similar to Du et al., 2024). For functional connectivity analyses we removed the first 25 TRs (40s) of every fixation run, leaving 260 sec (4.25 min) of data, to remove artifacts associated with the instructed deep breath during the second fixation run in each session. After these exclusions, one participant (subj03) had only a single fixation run remaining (4.25 min), and so was excluded from the present study’s analyses. The other 7 participants had between 8 and 24 runs (mean: 14, sd: 6.13, 34 – 102 min) of high-quality fixation data.

Retinotopy and fixation data were minimally preprocessed, including unwarping and aligning, as described in the NSD data release (Allen et al., 2022). We further denoised the data via manual ICA classification (Griffanti et al., 2014, 2017; Steel et al., 2021b), as automated ICA denoising tools have poor performance on high resolution data like the NSD (Beckers et al., 2023). For each run, the time series was decomposed into independent spatial components along with their associated temporal signal using FSL’s melodic ICA (Beckmann and Smith, 2004). We then assigned each component to either signal or noise using established criteria (Griffanti et al., 2017). After classification, all the noise components were regressed from the data (fsl_regfit; Griffanti et al., 2014), and the resulting denoised time course was normalized to percent signal change. We did not perform global signal regression (Saad et al., 2012), and all functional data was left unsmoothed.

### Retinotopic Mapping

#### Population Receptive Field Modeling

To perform population receptive field (pRF) modeling, we average all three denoised and normalized retinotopy runs together to produce an average retinotopy time series, to which we applied the AFNI modeling procedure using 3dNLfim as described in Silson et al. (2015). The model calculated 4 million possible 2D Gaussian receptive fields with a given center position and size based on the timing of the presented sweeping bar (resampled to the temporal resolution of the fMRI acquisition, TR = 1.333s). These ideal Gaussian responses were convolved with an a priori hemodynamic response function chosen to mirror a previous characterization of hippocampal activity during a visual oddball task (Angeli et al., 2025; peak activity at 4 s, FWHM ≈ 3.33 s). For each voxel, the best of these predicted time series was chosen after the application of a flexible amplitude scalar, using a combination of Simplex and Powell optimization algorithms, minimizing the least-squares error between the acquired BOLD time series and the amplitude-scaled prediction. Importantly, amplitude here reflects the relative activation/deactivation of a given voxel to visual stimulation of its receptive field. Group data was generated for visualization only by averaging the amplitude estimate across participants for all vertices on the AFNI standard 141 surface, without screening for significance (see Reliability and Stability of Hippocampal pRF Modeling below).

#### Reliability and Stability of pRF Modeling

To establish the reliability of our pRF model in the hippocampus, we compared task-derived model fits to those of a control null distribution for each participant. To generate this control null distribution of model fits, we applied the retinotopic model to hippocampal activity during a task that would not be expected to have systematic retinotopic visual field stimulation: resting-state fixation. For each participant, we first averaged the time course of all resting-state fixation runs prior to trimming, which were matched in length to the retinotopy task runs (300s). Then, we applied the retinotopic model to the average hippocampal resting state activity to derive model fits for each hippocampal voxel. Critically, because participants did not see the sweeping bar stimulus during resting-state fixation, this produces a null distribution of model parameters. We used this null distribution to determine an empirically-derived noise floor within the hippocampus. This was important because previous pRF research has applied R^2^ cutoffs based on data in visual areas, and it is not a given that the hippocampus would have a comparable noise-floor to visual areas. We chose a noise floor corresponding to the 95^th^ percentile R^2^ value of the null model (0.115), and only considered voxels with fits that exceeded this floor to be retinotopic. For consistency, and because null fits remained largely stable across regions (Fig. S4), reliability estimates in cortical regions were calculated using the same noise floor.

In addition to requiring fits above the noise floor, we conservatively screened voxels based on model estimates of size (sigma) and eccentricity, guided by the size of the presented stimulus (diameter 8.4°, bar width 1.05°). Specifically, we excluded any voxels with a size estimate smaller than .42° or larger than 3.78° (10% and 90% of the largest possible sigma estimate), as well as voxels with an eccentricity estimate greater than 3.78° (90% of maximum possible eccentricity). Voxels that passed the above noise floor, size, and eccentricity screens were determined to be “significant pRFs”, while all others were “non-significant voxels.” To avoid any bias resulting from exploring the same metric twice, we do not explicitly consider R^2^, size, or eccentricity in any of the subsequent analyses.

Finally, we sought to validate the stability of retinotopic features (amplitude sign and center position) between individual runs of the sweeping bar retinotopy task. To do so, we fit the above pRF model on each individual run of the sweeping bar stimulus, rather than the average, producing three sets of model parameters for each voxel. To ensure that voxel categorization would not differ across runs, we used model parameters derived from the three-run average to determine significance (based on model fit, size, and eccentricity as described above). To assess center position stability, we calculated the average Euclidean distance for each significant voxel between all three pairings of sweeping bar runs (i.e. run 1 – run 2, run 1 – run 3, and run 2 – run 3). We quantified amplitude sign stability by binarizing voxels according to their amplitude, and calculated a 3-run Dice coefficient 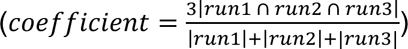. We applied this separately to positive amplitude voxels (i.e. significant pRFs with positive amplitudes binarized to 1, other significant pRFs set to 0) and negative amplitude voxels (significant pRFS with negative amplitudes binarized to 1, other significant pRFs set to 0), resulting in two, amplitude-specific, coefficients per participant. The significance of these stability measures was established on a participant-level by z-scoring the values described above (derived from significant voxels) against 5000 additional measures derived from a random sampling of all voxels, both significant and non-significant, matched in number to the number of significant pRFs.

#### ROI creation

To contextualize hippocampal pRFs within the ventral visual stream, we extracted cortical pRF data within three group-level ROIs: early visual cortex, the parahippocampal place area (PPA), and the ventral place memory area (vPMA). Early visual cortex was first defined on the participant-level to include V1, V2, and V3, as identified in the NSD release via retinotopic features (Allen et al., 2022). We then projected these participant specific ROIs to the standard 141 surface, and quantified the overlap across participants. Finally, the group ROI was constrained to vertices that fell within the early visual ROI in 4 or more participants. Group-level ROIs for PPA and vPMA were derived from publicly available dataset of probabilistic ROIs on the standard 141 surface (Steel et al., 2025), identified via scene perception tasks for PPA and scene memory tasks for vPMA. We included vertices with a probability of >20%, and further constrained the ROIs to be non-overlapping. This was achieved by visually identifying the anterior-posterior midpoint of the overlapping region, and defining PPA as posterior to that point, and vPMA as anterior. For consistency with the hippocampal analyses, pRF data was extracted in the volume by projecting each group ROI from the standard 141 surface into each participant’s volume-space.

Within the hippocampus, we defined ROIs for hippocampal subfields and relative position along the long (anterior-posterior) axis of the hippocampus. Participant-specific subfield ROIs, included in the NSD release, were calculated using the Freesurfer automated segmentation and visually inspected prior to use. To explore differences along the hippocampal long axis, each participant’s hippocampus was divided into 18 segments equally spaced along the y dimension of volume space.

#### Visual Field Coverage

To assess the visual field coverage of hippocampal pRFs, we applied a max-operator approach similar to Silson et al. (2021) and Steel et al. (2024b). In each participant, we reconstructed the 2D Gaussian corresponding to each significant pRF, with the amplitude scaled by that voxel’s model fit (R^2^). We refer to values derived from this Gaussian as pRF values. Next, for each participant we calculated the maximum pRF value at each point in the visual field across all significant pRFs with a given amplitude (positive or negative) and subsequently performed participant-specific normalization of pRF values such that the maximum value would be 1. These maximum-operator maps were then averaged to generate the group-level coverage maps. To calculate visual field bias, in each participant the X coordinate of peak maximum-value from the left hemisphere was averaged with the right hemisphere’s peak value X coordinate multiplied by −1. Statistical significance was assessed using a one-sided t-test with the alternative hypothesis that the mean visual field bias is greater than 0, indicating a contralateral bias.

#### Correlation of Anatomical Location and Visual Field Location

To explore the possibility that hippocampal retinotopy is organized in a sparse topographic map, we tested the relationship between hippocampal pRF’s preferred visual field location and that voxels distance from other significant hippocampal pRFs. To do so, we paired every hippocampal pRF with every other hippocampal pRF in the same hemisphere. We did both valence agnositic pairings (“All”) and valence-specific either within a valence (i.e. positive pRFs paired with only positive pRFs, and vice versa) or explicitly cross-valence (i.e. positive pRFs paired with only negative pRFs, and vice versa). For each pair, we then calculated the Euclidean distance between their pRF center positions and the Euclidian distance between their locations in the brain, and correlated those values across voxels within a participant.

### Functional Connectivity Analyses

#### Influence of Retinotopic Valence

We used a resting-state functional connectivity approach to explore how pRF model valence predicts intrahippocampal interactions. First, we averaged the time courses of hippocampal +pRFs, −pRFs, and non-significant voxels for each run of resting-state fixation. No spatial smoothing was applied to resting-state runs to preserve voxel-level differences, and keep preprocessing consistent with retinotopic mapping runs. We calculated the run-level partial correlation between the average + and −pRF time courses in each participant, controlling for global hippocampal correlation via the average non-significant voxel’s time course, then Fisher z-transformed the resulting correlations. Statistical significance was calculated at the participant-level via paired and one-sided t-tests.

#### Influence of Retinotopic Alignment

To better understand the influence of retinotopic features beyond valence on functional connectivity, we investigated differences in functional connectivity between pairs of pRFs sensitive to either similar or different portions of the visual field, which we term retinotopic alignment. To assess the influence of retinotopic alignment within the hippocampus, we took each hippocampal pRF sequentially as a source and sorted hippocampal pRFs within the same hemisphere of the same or opposite valence according to the Euclidian distance between that pRF’s center and the source pRF center. We defined the pRF with the smallest difference in Euclidian distance to be retinotopically aligned (“Matched”), and randomly sampled 1 voxel from the 1/3 of voxels with the largest distance 1000 times to generate a group of non-aligned voxels (“Random”). We then calculated the run-level partial correlation between the source voxel, the matched voxel, and each of the random voxels, controlling for the average activity of non-significant hippocampal voxels. Finally, these values were Fisher z-transformed, averaged across runs, and the random voxels further averaged to generate a single correlation value. After completing this process for every source voxel, we averaged the matched and random correlation values according to the valence of the source and target pRFs to generate 4 participant-level estimates for retinotopic alignment: 1) matched, same valence 2) matched, opposite valence 3) random, same valence 4) random, opposite valence.

### Statistical Tests

All statistical analyses were performed using R (version 4.3.2). Specific tests are described in the relevant portions of the Results and Methods. In summary: run-level significance for retinotopic modeling R^2^ was assessed using Kolmogorov-Smirnov test while other feature reliability measures (center position, valence) were z-scored relative to estimates sampled from all hippocampal voxels. We used a linear mixed effects model to assess the contribution of brain region to the proportion of −pRFs, controlling for mean R^2^. All other analyses were assessed using t-tests.

## Results

We first determined the presence of retinotopic coding in the human hippocampus. To do this, we performed voxel-scale population receptive field (pRF) modeling (Dumoulin and Wandell, 2008, for implementation see Cox, 1996, 2012) on high-resolution 7T fMRI data from the hippocampus during a visual retinotopic mapping task (Fig. 1A).

To validate the reliability of the task-driven retinotopic response in the hippocampus, we compared task-derived model fits to a control null distribution (Fig. 1B), which was generated by applying the retinotopic model to hippocampal activity during independent runs of resting-state fixation data from the same participants (see Methods). Task-derived model fits were significantly better than the null model fits (within participants: paired t-test, t(6) = 4.023, p < 0.01; across participants: asymptotic two-sample Kolmogorov-Smirnov Test, H_a_: task < null, D^−^ = 0.225, p < 0.001). To ensure that the hippocampal null distribution did not reflect an elevated baseline susceptibility to model noise relative to classic retinotopic regions, we compared resting-state null fits in the hippocampus to those in early visual cortex (EVC). Null R^2^ values were comparable across the regions (EVC R^2^ = 0.063 +/− 0.005; Hippocampus R^2^ = 0.060 +/− 0.002) (Fig. S4), indicating that the hippocampus is not uniquely susceptible to spurious retinotopic fits. We therefore defined a voxel as retinotopic if its pRF model fit exceeded the hippocampal noise floor, defined as the 95^th^ percentile of the null distribution (R^2^ = 0.115, Fig. 1B, see Methods). Voxels meeting these criteria were classified as retinotopic and are referred to throughout the manuscript as “pRFs” or “significant voxels”.

We observed pRFs throughout the hippocampus (Fig. 3A), with an approximately equal mix of positive and negative valences (+/− pRFs). We did not observe a significant hemispheric difference in the proportion or absolute number of significant voxels (proportion: t(6) = 0.243, *p* = 0.816; abs. number: t(6) = 0.211, *p* = 0.840), and so pooled analyses across both hemispheres. On average across all participants, approximately 9% (8.7%, +/− 1.70%) of hippocampal voxels were significantly retinotopic, with approximately half (51.7%, +/− 6.56%) of the significant voxels showing a negative valence (−pRFs). Negative pRFs, like positive pRFs, showed robust stimulus-evoked responses, characterized by stimulus-locked suppression of activity rather than increased activation (Fig. 1A, right). Importantly, positive and negative pRFs exhibited comparable average model fits (Fig. 1C; −pRF mean R^2^ = 0.162 +/− 0.007, +pRF mean R^2^ = 0.165 +/− 0.006; within participants: t(6) = 0.330, *p* = 0.753; across participants: D = 0.06, *p* = 0.658), and –pRFs were not preferentially associated with low R^2^ values prior to screening (Fig. S1), indicating that our modeling approach does not preferentially bias fits toward positive response amplitudes. Together, these findings indicate that hippocampal retinotopy is supported by both positive and negative response profiles that are equally robust and equally well captured by the pRF model.

As an additional validation, we assessed whether the retinotopic features of hippocampal pRFs were stable between identical runs of the visual mapping task. Hippocampal pRFs showed strong reliability across identical runs (Fig. 2A). Across pRF voxels, both center position estimates (i.e., preferred visual field location) and response valence (i.e., positive vs. negative responses) were highly reliable across different runs of the sweeping bar task, significantly outperforming a reference distribution of voxels generated by sampling from all hippocampal voxels (see Methods; all center position *p*-values < 0.001, avg. cross-run Euclidean distance, significant pRFs: 0.838±0.075, sampled voxels: 1.05±0.038; all positive valence *p*-values < 0.05, avg dice coefficient, significant pRFs: 0.544±0.129, sampled voxels: 0.295±0.068; 6/7 negative valence *p*-values < 0.05, avg dice coefficient, significant pRFs: 0.510±0.141, sampled pRFs: 0.256±0.066). While all reliability values decreased ascending the ventral visual stream from early visual cortex, the hippocampus showed comparable reliability to the nearby cortical ventral place memory area (vPMA) (Fig. S4B and C). Thus, hippocampal pRFs demonstrated run-level stability in both a core retinotopic property — preferred visual field location — and the more novel dimension of response valence.

**Fig. 2.**
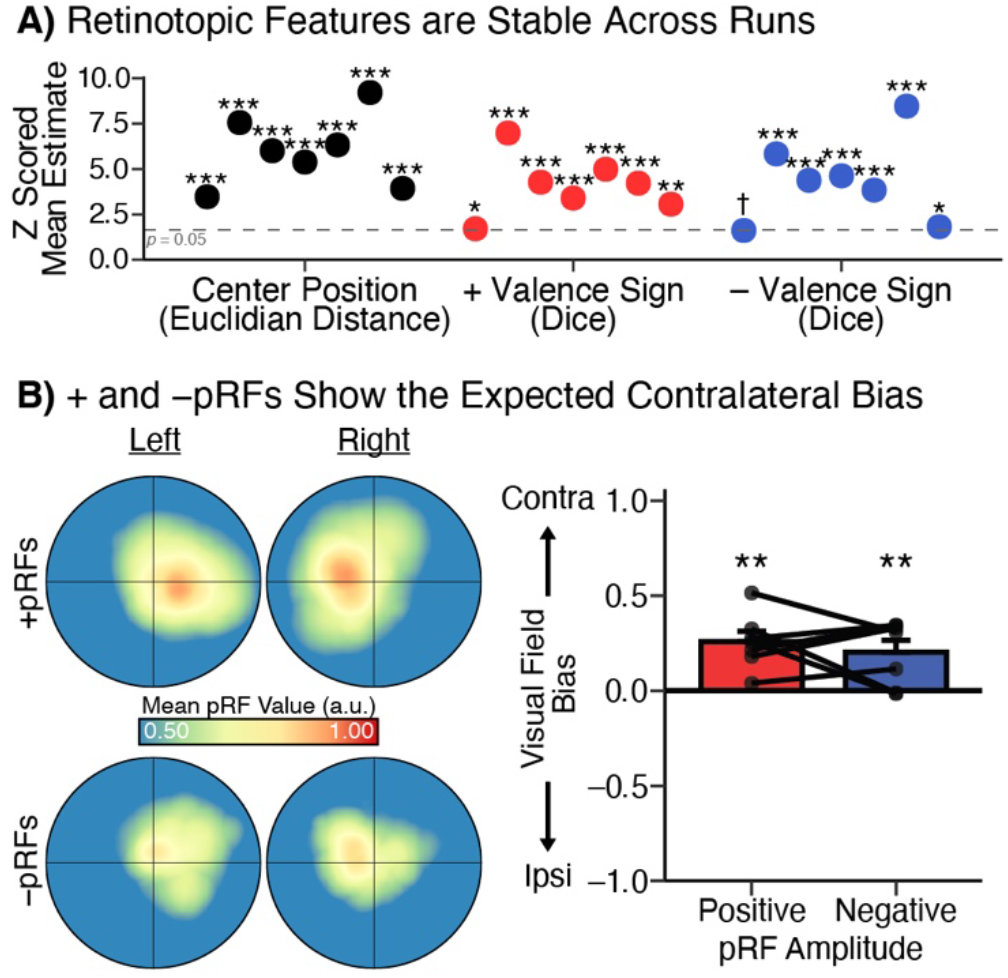
Hippocampal pRFs are stable across runs and exhibit contralateral bias. **A. Run-to-run stability of retinotopic features.** Center position (black) and valence (+, red; −, blue) were consistent across individual runs of the retinotopic mapping task. Center position consistency was calculated using average Euclidian distance between pairs of runs (black dots), while valence sign was calculated using a 3-run Dice coefficient (red and blue dots). Coefficients derived from voxels with significant pRF fits were z-scored relative to all hippocampal voxels. Each point is a participant, from left to right: subj01, subj02, subj04, subj05, subj06, subj07, and subj08. **B. Contralateral bias of + and − pRFs.** (Left) Mean visual field coverage using a max-operator (see Methods) (Right) Participant-level maximal bias shows that hippocampal +pRFs and –pRFs both preferentially cover the contralateral visual field (positive: 0.267 ± 0.05, one sided t-test, t(6) = 4.70, p < 0.01; negative: 0.2011 ± 0.068, t(6) = 3.20, p < 0.01). ***p < 0.001 **p < 0.01 *p < 0.05

Motivated by prior reports of a contralateral visual field bias in positive hippocampal pRFs (Knapen, 2021; Silson et al., 2021), we tested whether both valences of the observed bivalent code have the same bias. At the group level, both +pRFs and −pRFs demonstrated a significant bias towards the contralateral visual field (see Methods; +pRF mean bias: 0.26 +/− 0.05, t(6) = 4.70, *p* < 0.01; −pRF mean bias: 0.20 +/− 0.06, t(6) = 3.20, *p* < 0.01; Fig. 2B).

Extensive evidence suggests functional and organizational differentiation along the hippocampal long axis and subfields (for review see Poppenk et al., 2013; Strange et al., 2014); therefore, we next explored the relative concentration of +/− pRFs along the hippocampal long axis. We found a uniform spatial distribution of retinotopic voxels within the hippocampus (Fig. 3, see Table S1 for raw voxel counts), with no significant difference in the prevalence of significant pRFs along the long axis (Fig. 3B; F(1, 118) = 0.144, *p =* 0.705) or between subfields (Fig. 3C; F(8, 48) = 1.240, *p* = 0.297). This even distribution held when considering only +pRFs (long axis: F(1, 118) = 1.948, *p* = 0.165; subfields: F(8, 48) = 1.530, *p* = 0.172) or −pRFs (long axis: F(1, 118) = 2.01, *p* = 0.159; subfields: F(8, 48) = 1.322, *p* = 0.256). Additionally, we found long axis position was only a significant predictor of pRF size for –pRFs, with generally smaller size in the anterior hippocampus (Fig. S2). However, we interpret this size result with caution due to the limited extent of the mapping stimulus (8.4 d.v.a.). Despite this overall uniformity, we observed qualitative variability in the absolute prevalence of retinotopic voxels across subfields. The subiculum, CA4, and molecular and granular cell layers of the dentate gyrus showed the lowest overall proportion of retinotopic voxels (subiculum: 6.0% +/− 1.7%; CA4: 6.4% +/− 2.0%; DG Mol. and Gran.: 6.6% +/− 2.0%). Interpretation of the relative proportions of positive and negative pRFs within individual subfields is limited by the fact that some participants lacked significant pRFs in certain subregions, resulting in an uneven number of observations across subfields (see Table S1, n = 5 – 7). With this caveat, the parasubiculum has the highest prevalence of −pRFs (proportion of all pRFs: 70.0% +/− 18.7%, n = 5; proportion of all voxels: 9.3% +/− 5.0%, n = 7) while the molecular layer has the highest proportion of +pRFs (proportion of all pRFs: 61.8% +/− 12.3%, n = 7; proportion of all voxels: 3.0% +/− 1.6%, n = 7). Taken together, the broadly consistent prevalence of positive and negative pRFs across the hippocampus suggests that voxel-scale bivalent retinotopic organization is not restricted to a specific hippocampal subregion, while leaving open the possibility of finer-grained differences within subfields.

**Fig. 3.**
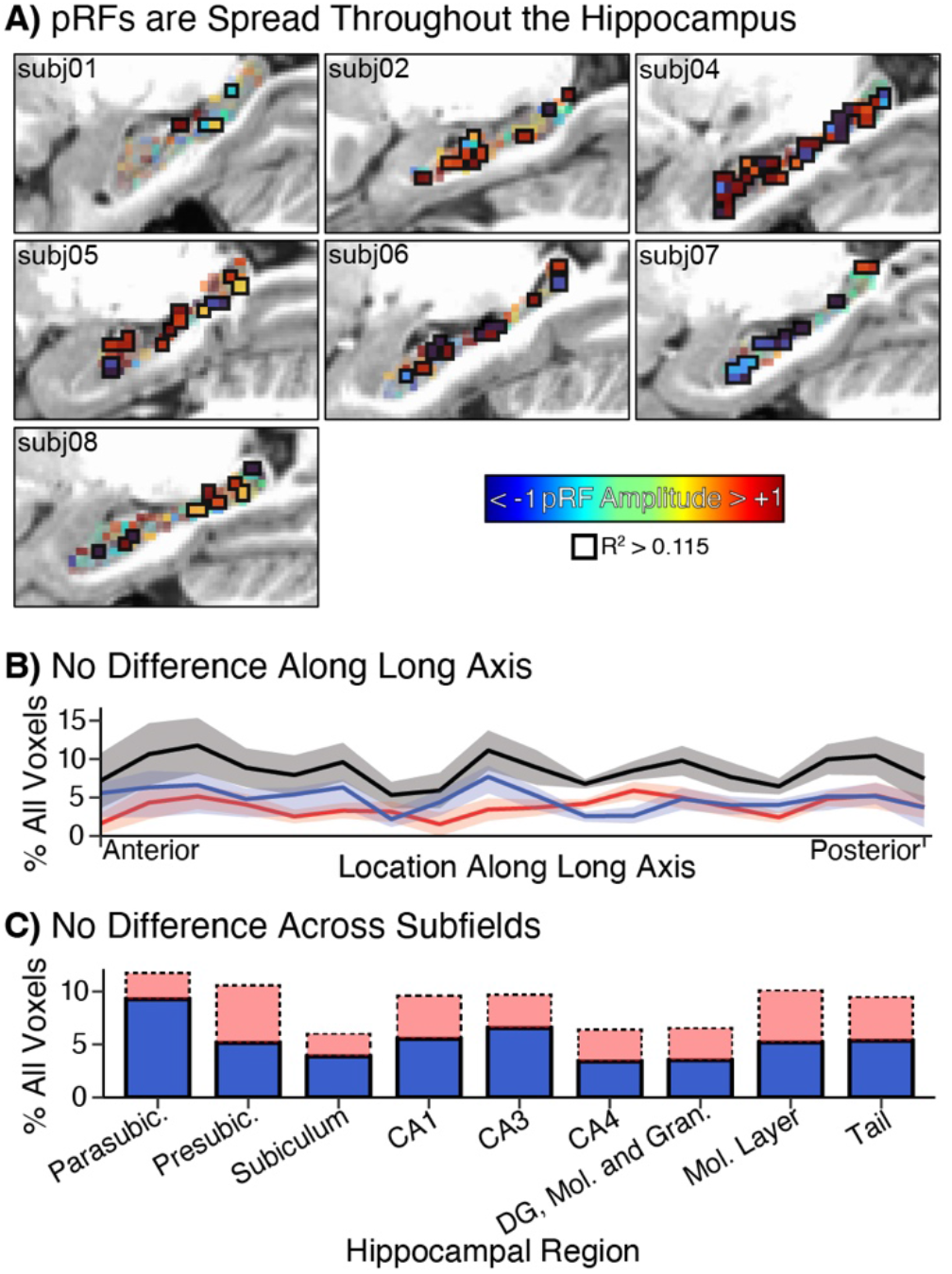
Retinotopic voxels are evenly distributed within the hippocampus. **A. Every participant displayed both positive and negative hippocampal pRFs.** Across participants, 9% of hippocampal voxels displayed significant pRFs. Voxel opacity reflects magnitude of R^2^ value, and voxels passing significance threshold of 0.115 are outlined. **B. No difference in pRF distribution along the long axis.** The proportion of significant pRFs (black line) and the proportion of each pRF amplitude (positive: red, negative: blue) calculated in 18 bins along the hippocampal long axis. Neither the proportion of significant pRFs (black line) nor the proportion of +/− pRFs (red and blue lines, respectively) significantly differed along the long axis (p = 0.159 – 0.705). **C. No difference in pRF distribution between subfields.** Despite qualitative differences in some subfields, notably the subiculum, we found no statistically significant difference across subfields in the overall proportion of significant pRFs or prevalence of specific valence (p = 0.172 – 0.297).

While the above analyses highlight the uniform distribution of pRFs across anatomical landmarks, they do not address the possibility that pRF features may cluster in anatomical space. Notably, pRFs in early visual regions are arranged in a topographic map such that nearby voxels in anatomical space also have nearby center positions in visual space. To assess whether this is also true for hippocampal pRFs, we calculated for every pair of pRFs the distance between their locations in the visual field and their anatomical locations in the brain (Fig. S3). Across subjects, the average overall correlation between visual field location and anatomical location in the hippocampus was not significantly greater than 0 (t(6) = 1.17, *p* = 0.287, Fig. S3A). When split by pRF valence (Fig. S3B), same-valence pairings also did not show a significant positive or negative correlation, while cross-valence pairings (i.e. positive pRF with negative pRF) demonstrated a significant negative correlation between visual field location and anatomical location (t(6) = −2.69, *p* < 0.05). These results indicate that hippocampal retinotopic responses are not organized in map-like configuration where similarly responding voxels are anatomically near to each other.

To provide functional context for the hippocampal retinotopic code, we explored whether pRF prevalence and valence follows the pattern established by visual regions lower in the hierarchy — where −pRFs become more prevalent towards mnemonic regions at the cortical apex, in line with their putative functional role in mnemonic processing (Steel et al., 2021a, 2023, 2024b) (Fig. 4). In early visual cortex, 52.4% (se: 1.7%) of voxels are retinotopic and only 14.3% (se: 3.1%) of those have a negative valence. Along the ventromedial visual pathway to the hippocampus (Rolls, 2025), the proportion of retinotopic voxels decreases while the proportion of negative voxels increases towards a maximum of 51.7% (se: 6.6%) in the hippocampus (Fig. 4). Parahippocampal cortex is of particular interest because previous work has identified a functional split between the perceptual parahippocampal place area (PPA) and mnemonic ventral place memory area (vPMA), found in the posterior and anterior portions, respectively, of parahippocampal cortex (Steel et al. 2021a, 2023). Mirroring this functional divide, 17.4% of pRFs in the PPA have a negative amplitude (se: 5.9%), while the immediately anterior vPMA has a much higher proportion of −pRFs (28.9% +/− 4.2%). To ensure that the relatively high proportion of negative pRFs in higher-order regions and the hippocampus was not driven by changes in the average model fit across regions (see Fig. S4), we fit a linear mixed-effects model predicting −pRF prevalence from region and mean R². Only region identity predicted −pRF prevalence (region: F(3, 18.305) = 8.825, *p* < 0.001; mean R^2^: F(1, 22.831) = 0.193, *p* = 0.665).Thus, consistent with previous work in cortex (Szinte and Knapen, 2020; Steel et al., 2024b), these results indicate that the proportion of −pRFs increases ascending the perceptual-mnemonic hierarchy, independent of retinotopic model fit, supporting the interpretation that +/− pRFs differentially relate to perceptual and mnemonic processing.

**Fig. 4.**
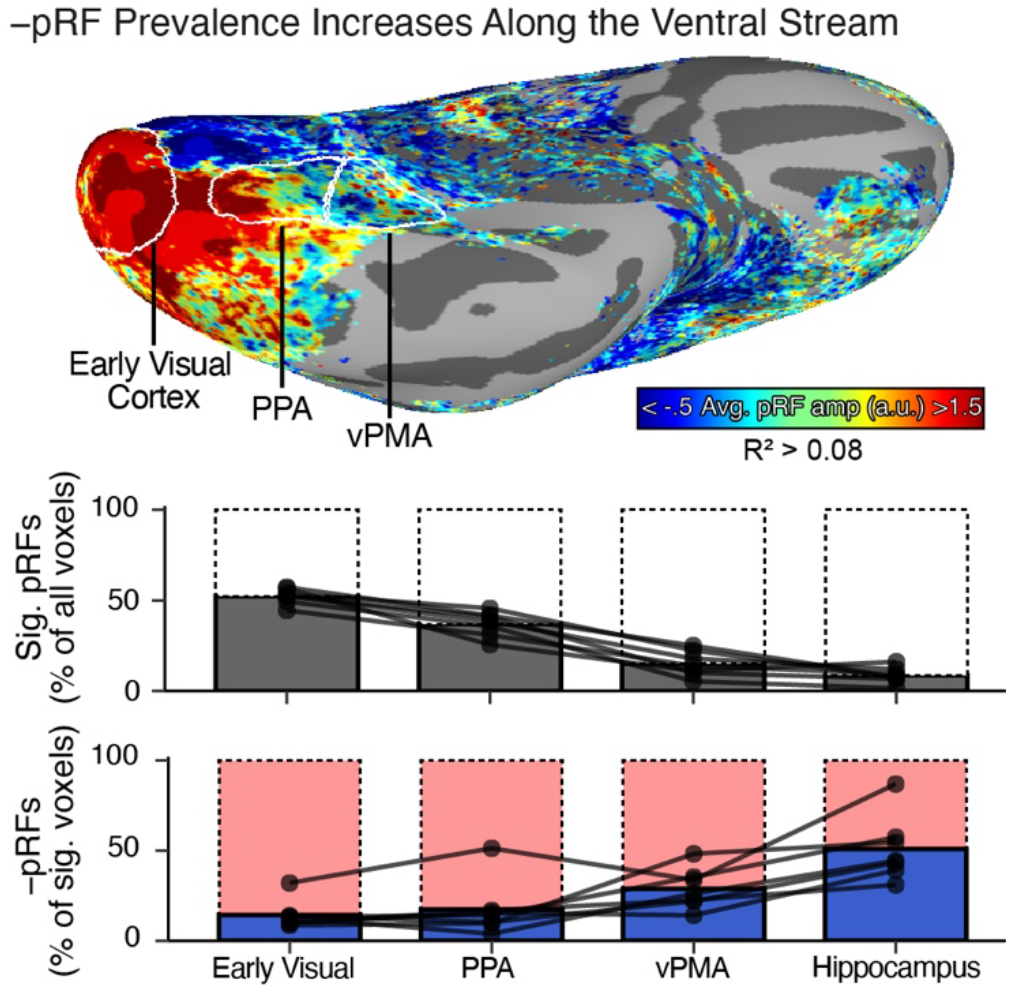
Negative pRF prevalence increases along the ventral visual stream. **(Top)** Cortical ROIs used for ventral stream hierarchy, overlayed on a group-average map of pRF amplitudes. Regions span visual to mnemonic areas along the ventromedial stream: early visual cortex (EVC); parahippocampal place area (PPA); and ventral place memory area (vPMA). **(Bottom)** Across participants, the overall proportion of significant retinotopic voxels (pRFs) is highest in visual areas and decreases in mnemonic areas. In contrast, −pRF prevalence increases moving up the visual-mnemonic hierarchy, culminating in the hippocampus, with the fewest pRFs and highest proportion of −pRFs.

In higher order cerebral cortex, including the Default Network (DN), bivalent retinotopic codes structure activity not only during perceptual-mnemonic tasks (Steel et al., 2024b), but also during resting-state fixation (Steel et al., 2024b), suggesting that retinotopy reflects an intrinsic organizational principle of spontaneous activity rather than a task or stimulus-driven effect. A defining feature of this architecture is antagonistic coupling between retinotopically-matched positive and negative pRF populations, thought to structure a push–pull interaction between externally oriented perceptual processing and internally oriented mnemonic processing (Steel et al., 2024a, 2024b). If the hippocampal bivalent code is similarly fundamental, it should continue to shape voxel-scale functional coupling even in the absence of structured visual input. We therefore tested whether the retinotopic properties of hippocampal voxels (valence, visual field location), as estimated during visual mapping, predict voxel-scale activity during independent resting-state fixation runs.

First, we tested whether the functional coupling of hippocampal pRFs during fixation was structured by their valence (i.e., positive vs. negative) (Fig. 5A). Critically, visual stimulation during fixation runs is limited to a small static plus sign in the center of the screen. Despite the absence of structured visual stimulation, activity in opposite-valence pRFs (i.e., positive correlated with negative) was generally anti-correlated at rest, after controlling for the activity of non-retinotopic voxels (z(r) = −0.166 +/− 0.071, t(6) = 2.35, *p* = 0.057). Though this anti-correlation did not reach statistical significance, it suggests the antagonistic functional relationship identified by bivalent retinotopy is not constrained to visual tasks.

**Fig. 5.**
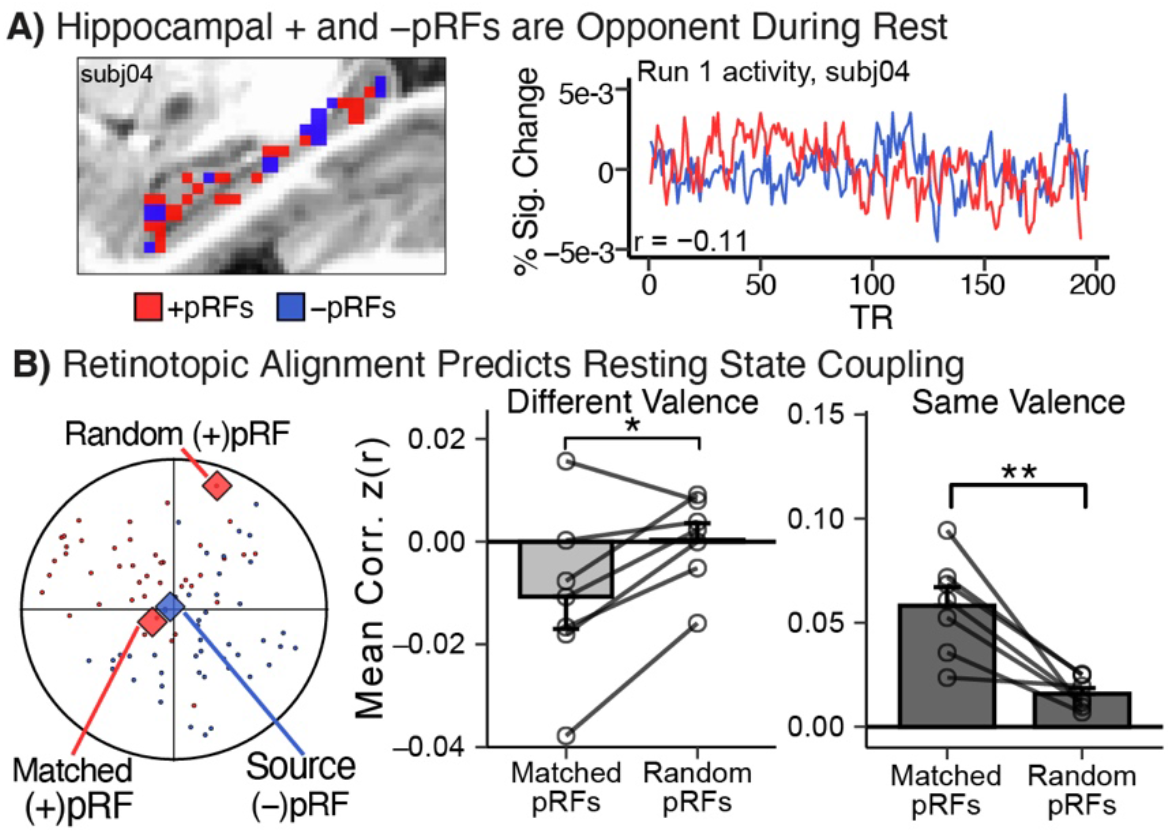
Retinotopic valence and alignment predict voxel-wise functional coupling within the hippocampus at rest. **A. Positive and negative pRFs are anti-correlated at rest (Left)** Example positive and negative hippocampal pRFs. (Right) Average positive (red) and negative (blue) pRF activity from an example participant during a single run of resting-state fixation, after regressing non-significant voxel activity. Note the broadly anticorrelated activity, despite the lack of dynamic visual stimulation. **B. Retinotopic alignment enhances pRF relationships at rest (Left)** Procedure for defining retinotopic alignment. Source pRF activity was correlated with the pRF with the closest visual field location (“Matched”), and this correlation was compared to the mean of a random sampling of pRFs from the furthest 1/3 of pRFs (“Random”). All correlations controlled for the activity of non-significant hippocampal voxels, and were performed for both same-valence (+/+; −/−) and opposite-valence (+/−) pRFs. **(Middle and Right)** Relative to randomly aligned pRFs, retinotopically aligned hippocampal pRFs of opposite valence were significantly more anti-correlated during fixation, while aligned pRFs of the same valence were significantly more positively correlated. Error bars indicate standard error around the mean. **p < 0.01 *p < 0.05

We next asked whether resting-state coupling was structured by retinotopic alignment — that is, whether voxels monitoring similar portions of visual space exhibited stronger functional relationships (Fig. 5B). Pairs of opposite-valence hippocampal pRFs (i.e. one positive and one negative pRF) that monitored nearby regions of the visual field (“Matched”) were significantly more anti-correlated than pairs of pRFs that monitored different locations of the visual field (“Random”, t(6) = 2.895, p < 0.05; see Methods). Conversely, same-valence pRFs with matched receptive fields were more positively correlated than random pairs (t(6) = 4.632, p < 0.01). Note that while the opposite-valence effect may be partially mediated by the negative correlation between visual field location and anatomical location for cross-valence pairings (Fig. S3B), this correlation does not exist for same-valence pRFs. Together, these results indicate that retinotopic features predict intrahippocampal functional connectivity at rest, consistent with the view that bivalent retinotopic mapping serves as a general scaffold for structuring spontaneous dynamics within the hippocampus.

## Discussion

Here, we identified a bivalent retinotopic code in the human hippocampus, with roughly equal proportions of positive and negative visually-evoked responses (+/−pRFs). These populations exhibit consistent valence and visual field preferences, show a contralateral bias, and are broadly distributed throughout the hippocampus. Critically, this organization extends beyond retinotopic mapping tasks: shared visual field preference – estimated during retinotopic mapping – continues to predict intrinsic functional coupling at rest. Together, these findings suggest that retinotopy in the hippocampus reflects an intrinsic organizational principle, rather than simply a consequence of visual stimulation, potentially serving to link perceptual and mnemonic processing at the apex of the visual hierarchy.

### Challenging Conventional Models of the Visual Hierarchy

Our findings challenge conventional views which propose that low-level sensory codes like retinotopy are shed as information ascends the visual hierarchy towards the hippocampus (Jeffery, 2007; Whitlock et al., 2008; Vann et al., 2009; Clark et al., 2018; Alexander et al., 2023). These models suggest that the hippocampus represents abstract, allocentric internal models of the external world at the expense of fine-grained sensory details present in early visual areas. However, our study joins previous work (Knapen, 2021; Silson et al., 2021) in demonstrating that low-level visual coding persists in the hippocampus, and further expands upon that work by characterizing the retinotopic code as bivalent in nature and persistent at rest.

Hippocampal pRFs remain constrained by classic sensory features such as contralateral visual field bias, as in visual (Hemond et al., 2007; Wandell et al., 2007; Chan et al., 2010; Groen et al., 2022a) and higher order cortex (Szinte and Knapen, 2020; Knapen, 2021; Silson et al., 2021; Steel et al., 2024b). This reinforces that visual field preferences are encoded in higher order, mnemonic cortex. Unlike in perceptual regions — where most retinotopic voxels are +pRFs — the hippocampus contains a balanced mixture of positive and negative retinotopic populations distributed throughout, with no topographic organization. Rather than reflecting a progressive transformation to purely mnemonic coding within the hippocampus, this bivalent organization indicates that perceptual and mnemonic representations are intermixed. This balanced architecture suggests a close linkage between the hippocampus and vision (Nau et al., 2018; Turk-Browne, 2019), while also underscoring the importance of negative retinotopic responses in mnemonic circuitry (Szinte and Knapen, 2020; Steel et al., 2024a, 2024b). Together, these findings support the view that bivalent retinotopy is a broader organizational principle in higher-order mnemonic regions, structuring persistent interactions with visual cortex.

### Integration with Other Hippocampal Neural Codes

The presence of a bivalent retinotopic code in the human hippocampus initially seems at odds with canonical descriptions of allocentric, spatial selectivity often described in hippocampal electrophysiology studies (O’Keefe and Dostrovsky, 1971; Moser et al., 2008). However, recent work suggests a complex, multi-faceted view of hippocampal activity that accommodates both response types — particularly in non-human primates (Piza et al., 2024). In freely moving macaques and marmosets, canonical place responses are rarely pure representations of spatial location (Gulli et al., 2020; Mao et al., 2021; Piza et al., 2024). Instead, hippocampal neurons most commonly present a multiplexed representation of allocentric space alongside egocentric features including head direction, facing location, or gaze.

Hippocampal egocentric visual codes are thought to be more prevalent in primates than in rodents due to differences in the spatial exploration strategy between species (Piza et al., 2024). Rodents are nocturnal, and supplement relatively coarse visual information with local whisking and olfaction while moving (Piza et al., 2024). In contrast, diurnal primates rely more on long-range visual information, with spatial exploration largely occurring while stationary (Piza et al., 2024). Yet even rodents display significant hippocampal mixed-selectivity to visual information (Acharya et al., 2016), with visual stimulation alone being sufficient to drive a subset of place cells in mice (Chen et al., 2013) and significant overlap (78%) between hippocampal neurons sensitive to egocentric vision and allocentric position (Purandare et al., 2022).

Within this broader framework of mixed selectivity, the relatively sparse retinotopic code we observe here (∼9% of hippocampal voxels) likely co-exists with spatiotopic codes of place and grid cells. Indeed, computational models of hippocampal dynamics suggest that place and grid cells could still serve as the dominant structure for information processing in the hippocampus, providing pre-structured states to support memory encoding in both spatial and non-spatial tasks, with sensory information applied to the scaffold of these pre-structured states (Chandra et al., 2025). Future research should explore whether the bivalent retinotopic code detected in this study is overlaid on top of place and grid cell representations, or whether it constitutes an independent contributing code that interacts with these spatial representations.

### Functional Relevance of Bivalent Hippocampal Retinotopy

Recent work in cortex suggests that bivalent retinotopy at the apex of the cortical hierarchy structures a push-pull dynamic between perceptual and mnemonic systems (Steel et al., 2024b, 2024a). We observe a similar oppositional relationship within the hippocampus, where pRFs of opposite valence are anti-correlated. Critically, the present findings demonstrate this organization within a single mnemonic structure, rather than across specialized regions (Steel et al., 2024b) or networks (Steel et al., 2024a), For example, a recent study found the externally-oriented Dorsal Attention Network and internally-oriented Default Network, classically considered functionally independent, show retinotopically-structured resting-state interactions (Steel et al., 2024a). The hippocampus appears to recapitulate this antagonistic logic at a local, intraregional scale.

One possible functional role for a bivalent code is supporting the coordination of perceptual encoding and mnemonic retrieval. In this context, opposing retinotopic populations could provide a spatial mechanism for separating concurrent encoding and retrieval-related processes while preserving a shared visuospatial reference frame, akin to the proposed role of hippocampal theta (Hasselmo et al., 2002; Kunec et al., 2005; Hasselmo and Stern, 2014; Kerrén et al., 2018). Notably, eye-movements provide a compelling link between visual input, theta oscillations (Hoffman et al., 2013; Jutras et al., 2013; Doucet et al., 2020), and hippocampally-dependent behavior (Wynn et al., 2020b, 2020a; Favila and Aly, 2023; Setton et al., 2024). Visual input (via retinotopy and theta) may thus allow the hippocampus to coordinate perceptual and mnemonic functions while maintaining functional separation.

Alternatively, the bivalent retinotopic code may reflect predictive coding processes (Steel et al., 2024b, Wirth, 2023). Negative pRFs could encode top-down sensory predictions, whereas positive pRFs reflect bottom-up sensory drive. Their antagonistic interaction would enable rapid detection of mismatches between expected and observed input, thereby supporting mnemonic updating. Future work should investigate whether valence-specific populations differentially track prediction errors or encode predictive templates during memory-guided perception.

### Open Questions and Future Directions

Several open questions lie beyond the scope of the present study. While we have shown that a pRF model robustly fits hippocampal responses during retinotopic mapping, these responses may not be fundamentally retinotopic in nature. Notably, work describing hippocampal view-selective cells (Rolls and O’Mara, 1995; Liu et al., 2017; Corrigan et al., 2023; Piza et al., 2024) suggests an alternative spatiotopic explanation in which hippocampal responses are linked to locations within a visual view, rather than locations of stimulation on the retina. This study is unable to disentangle these possibilities because participants were instructed to maintain fixation. Similarly, the temporal continuity of the moving bar stimulus makes it difficult to assess the potential impact of predictive responses in visually-evoked hippocampal activity.

The present data also leave unresolved questions about temporal dynamics, as the temporal resolution of fMRI (1.33s) precludes precise characterization of timing. As noted above, the hippocampal theta oscillation is implicated in sensory-mnemonic processing (Hasselmo et al., 2002; Kunec et al., 2005; Hoffman et al., 2013; Jutras et al., 2013; Hasselmo and Stern, 2014; Doucet et al., 2020), and bivalent retinotopy may be shaped by oscillatory dynamics. Further studies exploring hippocampal retinotopic responses with greater temporal resolution, such as with MEG or intracranial recordings, are needed to resolve this interaction and achieve descriptive parity with early visual retinotopic codes (Groen et al., 2022b; Yuasa et al., 2023).

Finally, there are limitations to the current work stemming from the type and quantity of collected tasks. First, only three runs of the sweeping bar task were acquired for each participant, and more data would likely improve hippocampal model fits. Second, because only resting-state fixation lacked dynamic visual stimulation, we used it for two analyses, creating some apparent tension. Rest was first used to define a non-visual null model floor, and later used for testing whether a retinotopic voxel’s amplitude and center position predicted intrinsic functional coupling. Because these properties were estimated from retinotopic mapping, rather than rest, this dual use should not bias our results. Nevertheless, future studies should consider using distinct non-visual tasks for these purposes.

To summarize, we identified a bivalent retinotopic code in the human hippocampus, which structures voxel-wise interactions within this mnemonic structure. This bivalent retinotopic scaffold may allow for the segregation and integration of perceptual and mnemonic information. By demonstrating that egocentric visual field coding persists within hippocampal circuitry, these findings expand the known extent of retinotopic organization in the human brain and establish a framework for investigating how this code contributes to memory-guided behavior.

## Data availability

All raw and minimally processed data is made publicly available via the Natural Scenes Dataset (naturalscenesdataset.org). All code required to reproduce the results in this manuscript will be made available in the Open Science Framework repository upon publication.

## Acknowledgements

The authors would like to thank the Natural Scene Dataset authors for making their data publicly available. This work was supported by funding from the National Institutes of Mental Health under award number R01MH130529 to CER.

## Author contributions

P.A.A., A.S., and C.E.R designed the study, analyzed the data, and interpreted the experiments; P.A.A. and C.E.R. wrote and edited the manuscript; A.S. edited the manuscript. C.E.R. secured funding.

## Supplemental Figures

**Figure S1.**
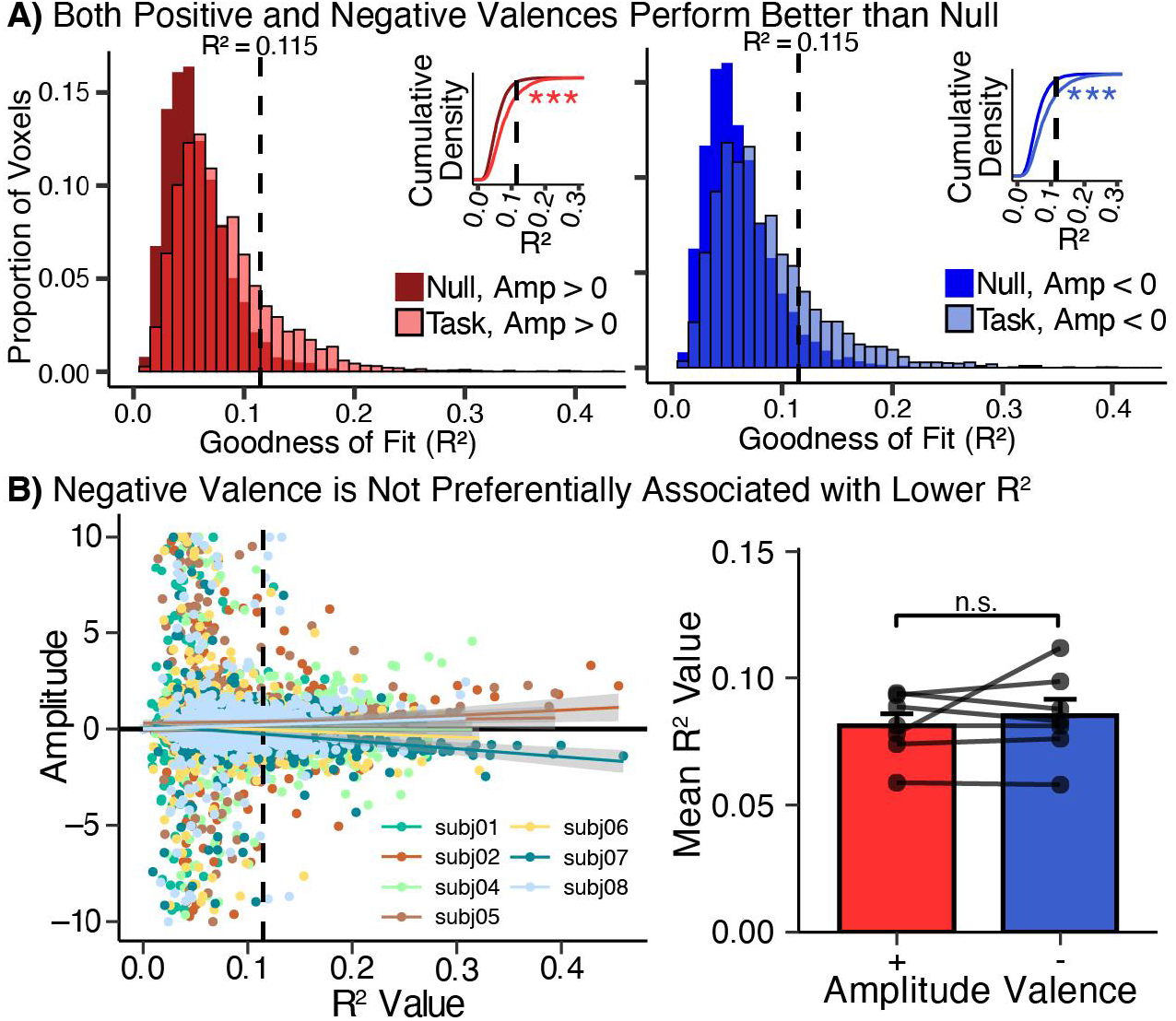
Positive and negative valences are not differentially associated with model fit within the hippocampus. A) Comparison of Null and Task model fits restricted to either positive (red) or negative amplitude (blue) fits, as in Figure 1B. Both valences have significantly better model fits to the Task data relative to the Null data. B) Both continuous amplitude (left) and binarized amplitude valence (right) are not significantly associated with model R^2^. For continuous amplitude, 5/7 participants showed no significant relationship between R^2^ and amplitude (p = 0.099 - 0.907). Subj02 showed a significant positive association (_β_ = 2.69, p < 0.01), while subj07 showed a significant negative relationship (_β_ = −4.13, p < 0.001). The average R^2^ _β_-estimate was not significantly different from 0 (t(6) = −0.008, p = 0.994). Binarized amplitude valence was not significantly different between positive and negative valence (t(6) = −0.777, p = 0.467).

**Figure S2.**
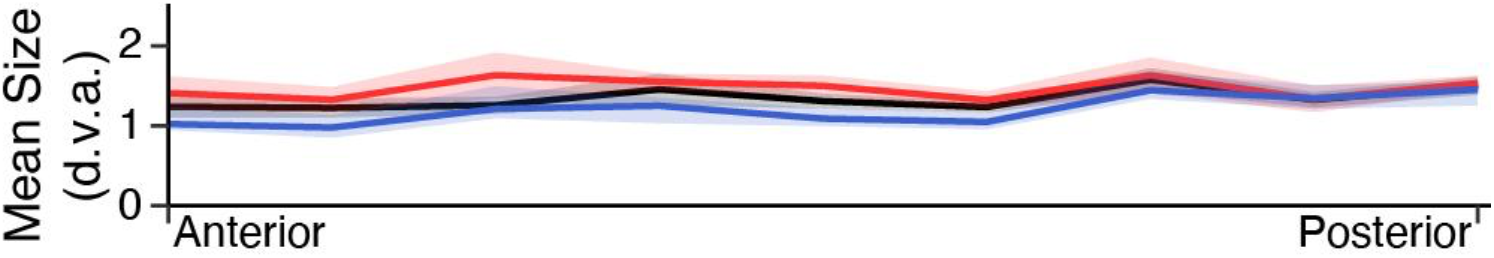
Changes in mean pRF size along the long axis. Positive pRFs (red) and the mean of all pRFs (black) do not show a significant difference in size along the long axis. Negative pRF sizes (blue) are significantly predicted by anterior-posterior location, with smaller pRFs in the anterior (F(1,53) = 9.4246, p < 0.01).

**Figure S3.**
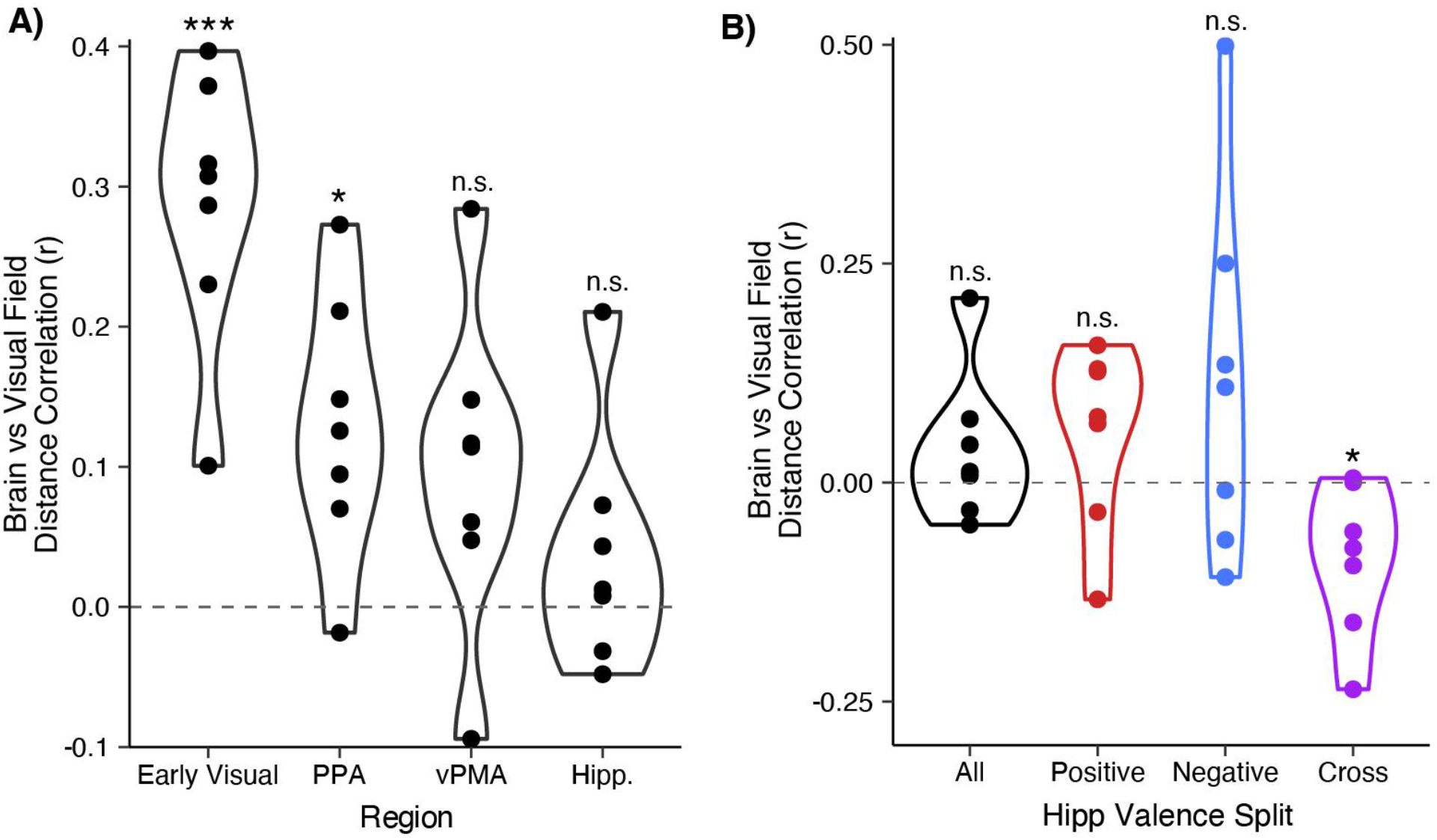
Relationship between brain location and preferred visual field location for significant pRFs. A. Visual regions demonstrate a significant correlation between brain location and visual field location, while mnemonic regions (vPMA, hippocampus) do not. B. Within the hippocampus, only cross-valence pairings (e.g. positive pRF with negative pRF) show a significant correlation between brain and visual field distance. * p < 0.05, ** p < 0.01

**Figure S4.**
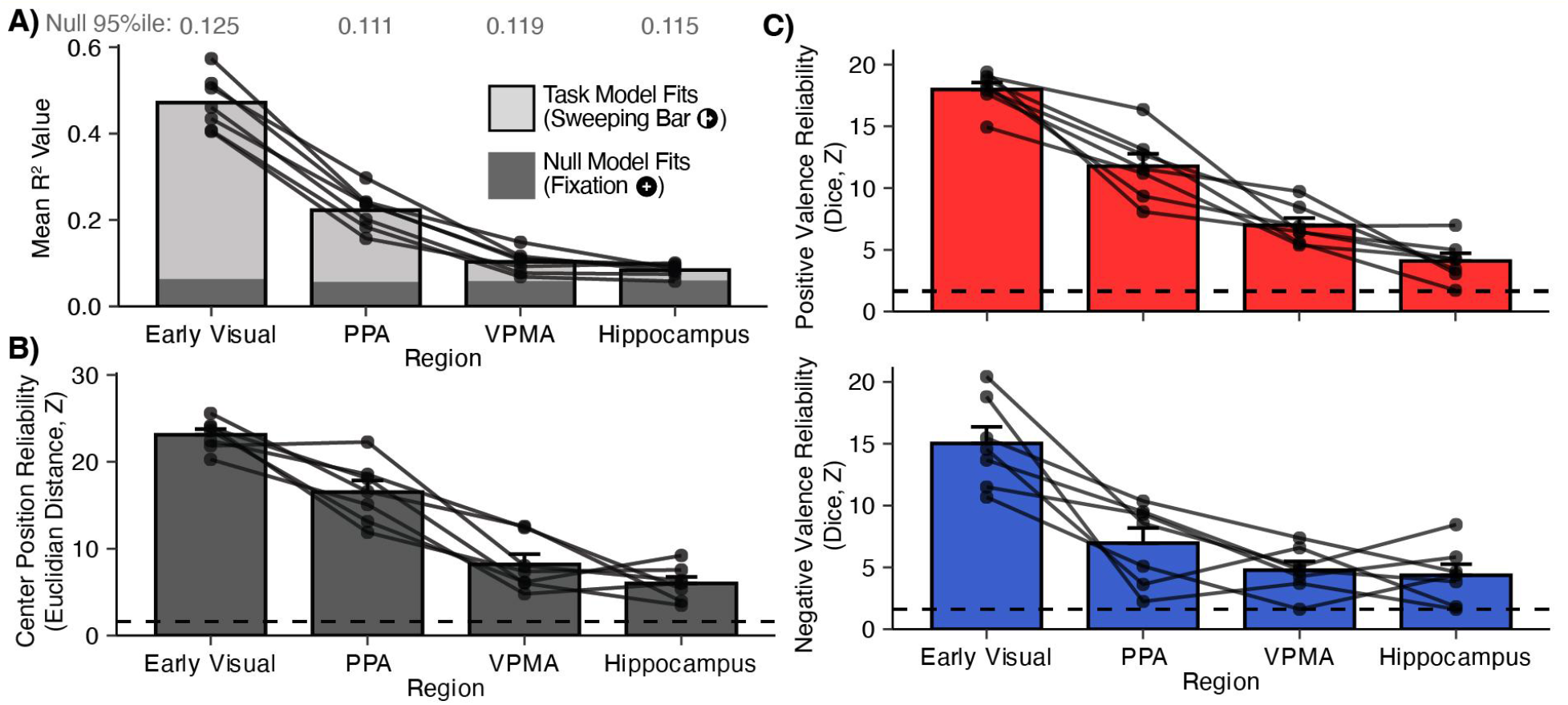
Retinotopic model fit and reliability along the ventral visual stream. A. Moving along the visual hierarchy reduces the average model fit, but does not influence the noise floor derived from resting-state fixation. Mixed effects modeling shows that region, rather than mean R^2^ value, is the primary driver of –pRF proportion (region: F(3, 18.305) = 8.825, p < 0.001; mean R^2^: F(1, 22.831) = 0.193, p = 0.665). Text above bars indicates the value of the 95th percentile R^2^ value from the null model for each region. B. The run-level reliability of center position estimates decreases moving up the hierarchy, but remains significant in participants to all regions. C. Valence reliability decreases moving up the hierarchy, but largely remains significant. Negative valence reliability was similar between the hippocampus and vPMA. Dashed line indicates z-value corresponding to p = 0.05.

